# Extraction-free direct PCR from dried serum spots permits HBV genotyping and RAS identification by Sanger and minION sequencing

**DOI:** 10.1101/552539

**Authors:** Stuart Astbury, Marcia Maria Da Costa Nunes Soares, Emmanuel Peprah, Barnabas King, Ana Carolina Gomes Jardim, Jacqueline Farinha Shimizu, Paywast Jalal, Chiman H Saeed, Furat T Sabeer, William L Irving, Alexander W Tarr, C Patrick McClure

## Abstract

In order to achieve the commitment made by the World Health Organisation to eliminate viral hepatitis by 2030, it is essential that clinicians can obtain basic sequencing data for hepatitis B virus (HBV) infected patients. While accurate diagnosis of HBV is achievable in most clinical settings, genotyping and identification of resistance-associated substitutions (RAS) present a practical challenge in regions with limited healthcare and biotechnology infrastructure. Here we outline two workflows for generating clinically relevant HBV sequence data directly from dried serum spot (DSS) cards without DNA extraction using either Sanger, or the portable MinION sequencing platforms. Data obtained from the two platforms were highly consistent and allowed determination of HBV genotype and RAS. This is the first demonstration of MinION sequencing from DSS, illustrating the broad utility of this sequencing technology. We demonstrated the clinical application of this technology using sera sampled on DSS cards obtained from both Iraq and Brazil. The sample stability provided by DSS cards, combined with the rapid PCR and sequencing protocols will enable regional/national centres to provide information relevant to patient management. By providing viable workflows for both the Sanger and MinION sequencing platforms, which vary greatly in the infrastructure and expertise required, we demonstrate that MinION sequencing is a viable method for HBV genotyping in resource-limited settings. These workflows could also be applied to sequencing of other blood borne DNA viruses and bacterial pathogens.

## Introduction

Hepatitis B virus (HBV) infects 257 million people worldwide and causes 887,000 deaths per year, many due to hepatocellular carcinoma (HCC) (1). A recent editorial in *Nature* described HBV as neglected, particularly in Sub-Saharan Africa, where around 6% of the population are infected and only one-tenth of children are vaccinated (2). There is an urgent need for screening and surveillance tools to assess HBV in low and middle-income countries (3). HBV displays a higher degree of diversity relative to other DNA viruses due to a complex and error prone replication cycle involving RNA intermediates, as such it has a mutation rate of around 1.4-3.2 × 10^-5^ mutations/site/year (4). There are many well-characterised mutations across the HBV genome, conferring resistance to therapy or an increase in replication efficiency (polymerase), immune and diagnostic test escape (S/pre-core), or an increase in tumorigenesis (reviewed in (5)).

Next generation sequencing (NGS) enables the study of a population of viral genomes within a single patient, as opposed to Sanger sequencing, which will theoretically provide a “consensus” sequence of the most abundant template. NGS platforms such as those established by Illumina rely on the use of short reads (max 250bp) which are assembled to from a contiguous sequence. Third generation sequencing platforms such as the Oxford Nanopore (ONT) MinION enable sequencing with theoretically no upper limit on read length, meaning entire viral genes or genomes can be sequenced in a single read. The MinION sequencer is also extremely portable and can be powered through a laptop, removing the need for a continuous power supply and enabling sequencing in locations without access to conventional lab facilities. The portable aspect of MinION sequencing has already been demonstrated to great effect during the Ebola outbreak in West Africa (6), and in tracking the spread of Zika virus in Brazil (7). Field application of the MinION platform has been enhanced with recent advances including improved “R9.4” flow cells with increased accuracy and software such as Nanopolish (8), which works with signal-level data from the sequencer allowing generation of more accurate consensus sequences. However, despite the emerging potential of NGS technologies, Sanger sequencing remains the gold standard in clinical applications and represents the most accessible and affordable choice globally.

We thus aimed to combine both Sanger and MinION sequencing with dried serum spot (DSS) sampling, to develop a method by which HBV can be rapidly sequenced for both genotype as well as potential resistance-associated substitutions (RAS) that can be applied in regions without access to conventional sample storage or a cold chain. As this assay targets the overlapping ORFs containing S and the reverse transcriptase domain of the polymerase gene it can be used to genotype and detect resistance to antiretroviral treatment. HBV genomes are particularly well suited to direct PCR, enabling a clinical sample to be added directly to a PCR reaction with no extraction steps. However, with minor alterations this method can be applied to a wide range of viruses or bacterial pathogens.

## Methods

### Samples

Thirty HBV DNA-positive serum samples, with known viral load, were obtained from eight specialized centres in São Paulo State, Brazil, collected between July 2016 and April 2017. Samples varied in virus titre from 6.35 x10^2^ to 8.16 × 10^8^ international units (IU)/mL. In addition, 70 serum samples identified as HBV DNA-positive were collected and processed at the Erbil Central Laboratory, Iraq, in 2017. Samples varied in virus titre from 22 to 9.4 × 10^8^ IU/mL. The initial cohort of samples for sequence determination and primer optimisation were obtained from Nottingham University Hospitals NHS Trust, Nottingham, UK, and were held under ethical approval of the Nottingham Health Science Biobank. All samples were obtained for routine diagnostic investigation and surplus material was used in this study. Serum was stored at −70 °C prior to direct use or spotting in Nottingham and −20 °C in Brazil and Iraq.

### Viral load determination

Viral load in the Brazilian samples was determined by RealTime HBV Amplification Kit (Abbott). In Iraq, DNA was extracted from 200µl serum using the EZ1 Virus Mini Kit v2.0 (Qiagen), and viral load determined by Artus HBV PCR kit (Qiagen).

### Preparation of dried serum spot (DSS) cards

A volume of 25-30 µL serum was spotted onto a Whatman^®^ Protein Saver™ 903 Card (GE Healthcare), air-dried at room temperature for approximately 2 hours and stored at 4 °C until further use. 25 μL of serum typically saturated the 113.1 mm^2^ (12 mm diameter) area demarcated on the DSS cards, thus a 3mm diameter punch (7.07 mm^2^ area) represented approximately 1.5 µL of serum. Between each sample a clean 3mm^2^ punch was made to prevent sample carryover, and a clean filter paper control was included in each PCR run.

### PCR

HBV S gene-specific primers of HBV are shown in table 1. Two different modified polymerase enzymes were evaluated for this method: Hemo KlenTaq (NEB) and Phusion Blood Direct (Thermo Fisher). Hemo KlenTaq reactions were set up as 25 µL reactions containing 5 μL Hemo KlenTaq reaction buffer, 0.2 mM dNTP (Sigma), 2 µL Hemo KlenTaq enzyme, 0.3 µM of each primer and 2 µL serum. Phusion Blood Direct reactions were also set up as 25μL reactions containing 12.5 μL 2x Phusion Blood Direct mastermix, 0.5 μM of each primer, 8.5 μL water and 2μL of serum. DSS reactions were prepared in the same way, with a 3 mm^2^ punch used in place of serum and the volume adjusted to 25 µL with water. Cycling conditions were as follows, for Phusion Blood Direct: 98 °C for 5 minutes, 55 cycles of [98 °C for 1 second, 50 °C for 5 seconds and 72 °C for 20 seconds/kb], and a final extension at 72 °C for 1 minute. Cycling conditions for Hemo KlenTaq were: 95 °C for 3 minutes, 55 cycles of [95 °C for 20 seconds, 50 °C for 30 seconds and 68 °C for 2 minutes/kb], and a final extension at 68 °C for 10 minutes. The sensitivity of this assay with different primer pairs was also tested by serially diluting a high viral titre sample (1.2 × 10^7^ IU/mL) logarithmically down to 1.2 × 10^1^ IU/mL.

**Table 1:**
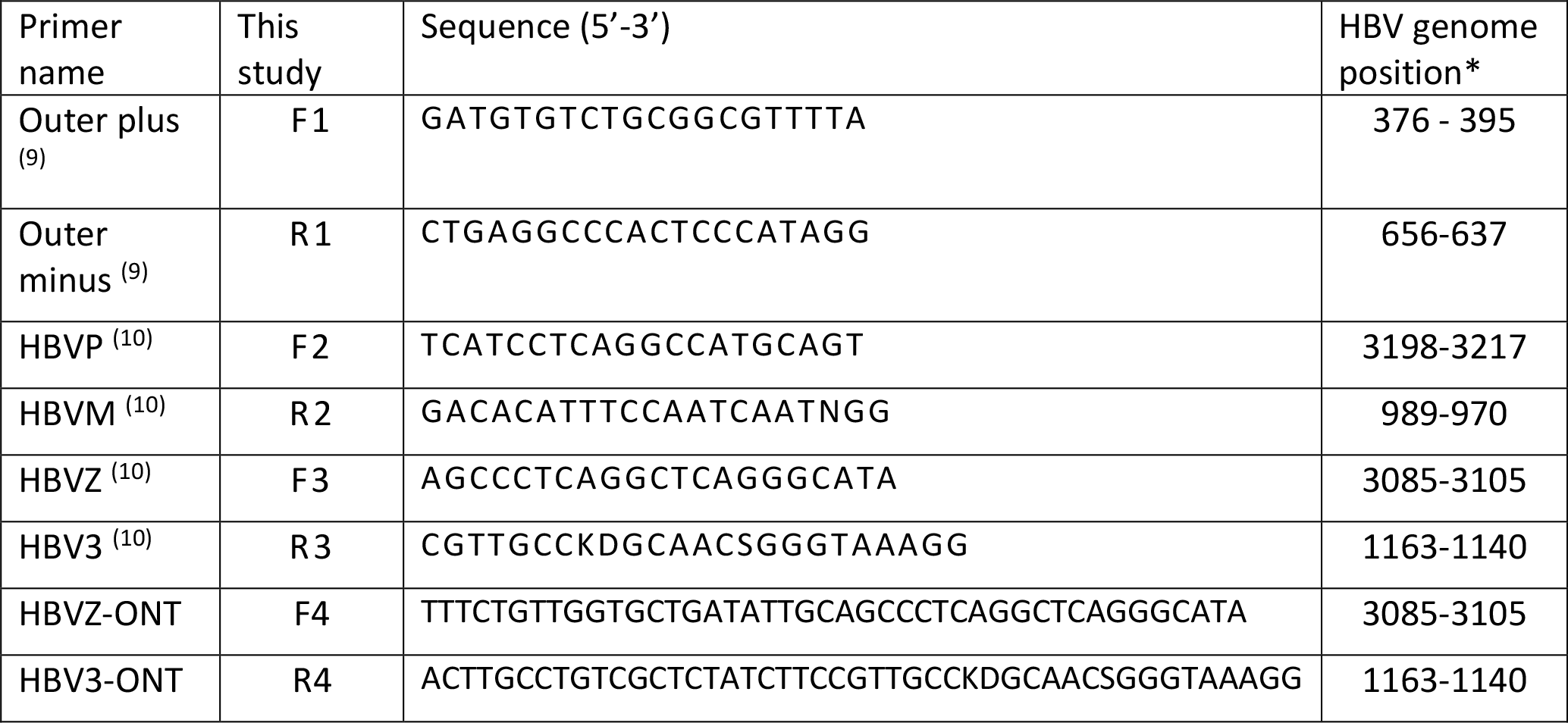
Primers used for the amplification of HBV. F4 and R4 primers contain 22 bp sequences at the 5’ end for MinION library preparation PCR. *Numbering based upon HBVdb genotype A master strain X02763 (17).

Initially a well cited primer set (F1 and R1) generating a short amplicon in a conserved genomic sequence towards the end of the S gene (9) was used to probe the maximum sensitivity of direct PCR from serum and DSS derived from the control Nottingham cohort. This primer site is also situated within the RT domain of the polymerase gene, allowing for genotyping and determination of some RAS. A second widely used primer set (F2, F3, R2 and R3; (10)) was subsequently tested to generate larger amplicons and thus greater DNA sequence information. The sets were further tested in a variety of combinations to generate amplicons of intermediate length overlapping in genomic coverage.

### Sanger sequencing and phylogeny

PCR-positive products were diluted 1:10 with water and couriered at ambient temperature for sequencing by a commercial service (Source Bioscience, Nottingham, UK). Short products were sequenced with F1 primer only, and longer F1/R3 and F3/R1 products were sequenced with both F1 and R1 primers. These reads were assembled to produce a complete contiguous sequence covering PreS2/S. Sequences used for phylogenetic analysis were aligned using MUSCLE in MEGA X (11, 12), and maximum likelihood trees constructed using a General Time Reversible model in MEGA X. Samples containing deletions were excluded from the tree. Sequences were deposited in GenBank under accession numbers MK517481 – MK517524.

### MinION sequencing library preparation

Three high titre (>10^7^ IU/mL) samples from both Brazil and Iraq cohorts were selected for sequence analysis. Barcoded samples were prepared for MinION sequencing using a two stage PCR approach and the SQK-LWB001 library preparation kit. First round PCR was carried out as above, using 25 µL Hemo KlenTaq reactions with 3 mm^2^ punches from DSS. F4 and R4 primers were used as specified in table 1. From this reaction, 1 µL of product was then used as the template for a 50 µl PCR containing 25 µL 2x LongAmp HotStart Taq master mix, 1.5 µL of barcoded primer mix (supplied by ONT) and 22.5 µL nuclease free water. Products were then prepared according to the SQK-LWB001 kit protocol. Briefly, PCR products were bound to AMPure XP beads (Beckman Coulter) and washed with 70 % ethanol, before eluting in 10 µL 10mM Tris-HCl (pH 8.0) with 50mM NaCl. Cleaned products were then quantified by Qubit using the dsDNA high sensitivity kit (Thermo Fisher). Based on the method developed by Quick *et al*. (7), amplicons were pooled to achieve a total of 0.3 pM input DNA per MinION flow cell, therefore for 1.3 kb amplicons this equates to ~260 ng total DNA. Following ligation of rapid sequencing adapters, the library was run for 48hrs on a MinION Mk II through a computer running MinKNOW 1.10.16 in Microsoft Windows followed by basecalling using Albacore 2.2.2. Adapter-trimmed sequences were uploaded to the NCBI Sequence Read Archive under project ID PRJNA521740.

### MinION sequencing analysis

Basecalled reads were trimmed using Porechop 0.2.3 using high stringency settings (--discard_middle and --require_two_barcodes) and retained when Porechop and Albacore barcode aligners were in agreement. NanoPlot was used to inspect read quality and length, and reads were filtered based on length using NanoFilt (13) to a minimum length of 1200 and maximum length of 1300 nucleotides.

Processed reads were used as the input for Canu (v1.7.1) (14), a *de novo* genome assembler optimised for use with long, error-prone reads. The same reads were subsequently aligned using Minimap2 (15) to their respective consensus sequences generated by Canu and manually inspected using IGV (Broad Institute) to ensure complete coverage and to inspect for significant structural variants. Alignments were further processed using Nanopolish v0.10.2 (8), using the Variants module and the --fix-homopolymers function to generate a corrected consensus sequence. A full description of the bioinformatics workflow used is included as supplementary data. As validation Sanger sequences were produced by sequencing amplicons with both F1 and R1 primers, these were used as the “gold standard”. Canu and Nanopolish consensus sequences were aligned with their respective Sanger contig using MEGA.

### Genotyping and variant calling

For both Sanger and MinION sequence data genotypes were determined using the web-based tool HBV geno2pheno (16). Known HBV treatment resistance and immune escape variants were screened using the hepatitis B virus database (HBVdb (17)). Mutations against the genotype consensus reference sequences used in geno2pheno that were not flagged as clinically significant are provided in supplementary table 1. For samples sequenced by MinION potential intra-host variants were screened for by aligning sequencing reads to the corresponding nanopolished consensus sequence for each sample and using the *variants* module within Nanopolish, with 0.1 set as the minimum frequency required to call a SNP.

## Results

### Primer analysis for PCR amplification of HBV from DSS

Initially we assessed three previously described primer sets for amplification of regions of the HBV genome using Hemo KlenTaq polymerase. A known HBV-positive serum sample with viral load >10^7^ IU/mL was diluted 10-fold and spotted onto DSS cards. Using direct PCR of punched-out discs from these cards, HBV amplicons were obtained using all the primer pairs analysed (table 2). Amplification of the smallest product, using primers F1 and R1, was the most sensitive approach, amplifying a product from a sample containing as little as 1 × 10^3^ IU/mL. Larger products (over 700 bp) were achieved using these primers in combination with the F3 and R3 primers down to 10^4^ IU/mL. Subsequent investigation demonstrated that PCR amplification using Phusion Blood Direct polymerase allowed for increased sensitivity when amplifying small products, down to 600 IU/mL but gave little improvement in sensitivity when amplifying larger products (data not shown).

**Table 2:** HBV amplification success using different primer combinations. HBV amplicon sizes at different viral loads from DSS using a serially diluted high-titre HBV-positive sample. Predicted amplicon size (in base pairs) is noted below primer pairing in brackets. +, indicates successful amplification or single discrete band following analysis by agarose gel electrophoresis.

| IU/mL HBV | Primer pair |  |  |  |  |  |
| --- | --- | --- | --- | --- | --- | --- |
|  | F1+R1<br>(281 bp) | F2+R2<br>(1,013 bp) | F1+R2<br>(614 bp) | F2+R1<br>(680 bp) | F1+R3<br>(788 bp) | F3+R1<br>(793 bp) |
| $1.2 \times 10^7$ | + | + | + | + | + | + |
| $1.2 \times 10^6$ | + | + | + | + | + | + |
| $1.2 \times 10^5$ | + | + | + | + | + | + |
| $1.2 \times 10^4$ | + | | + | + | + | + |
| $1.2 \times 10^3$ | + | | | | + | + |
| $1.2 \times 10^2$ | | | | | | |
| $1.2 \times 10^1$ | | | | | | |

### Sanger sequencing direct from DSS

Following this assessment, primer pairs F1/R1, F1/R3 and F3/R1 (figure 1) were used with Phusion Blood Direct polymerase for the analysis of 30 HBV-positive serum samples of defined viral load isolated in Brazil (table 3). Limited treatment information was available for some of the samples. Amplicons were achieved for all but 3 samples using the F1/R1 primer pair. The amplicon generated by this primer pair (281 bp) was sufficient to enable virus genotyping and limited detection of escape mutants by conventional Sanger sequencing. The 27 successfully genotyped samples divided into subtypes as follows: seven A1, two A2, one B1, two D1, two D1, two D2, nine D3, one F1 and three F2 (table 3).

**Figure 1.**
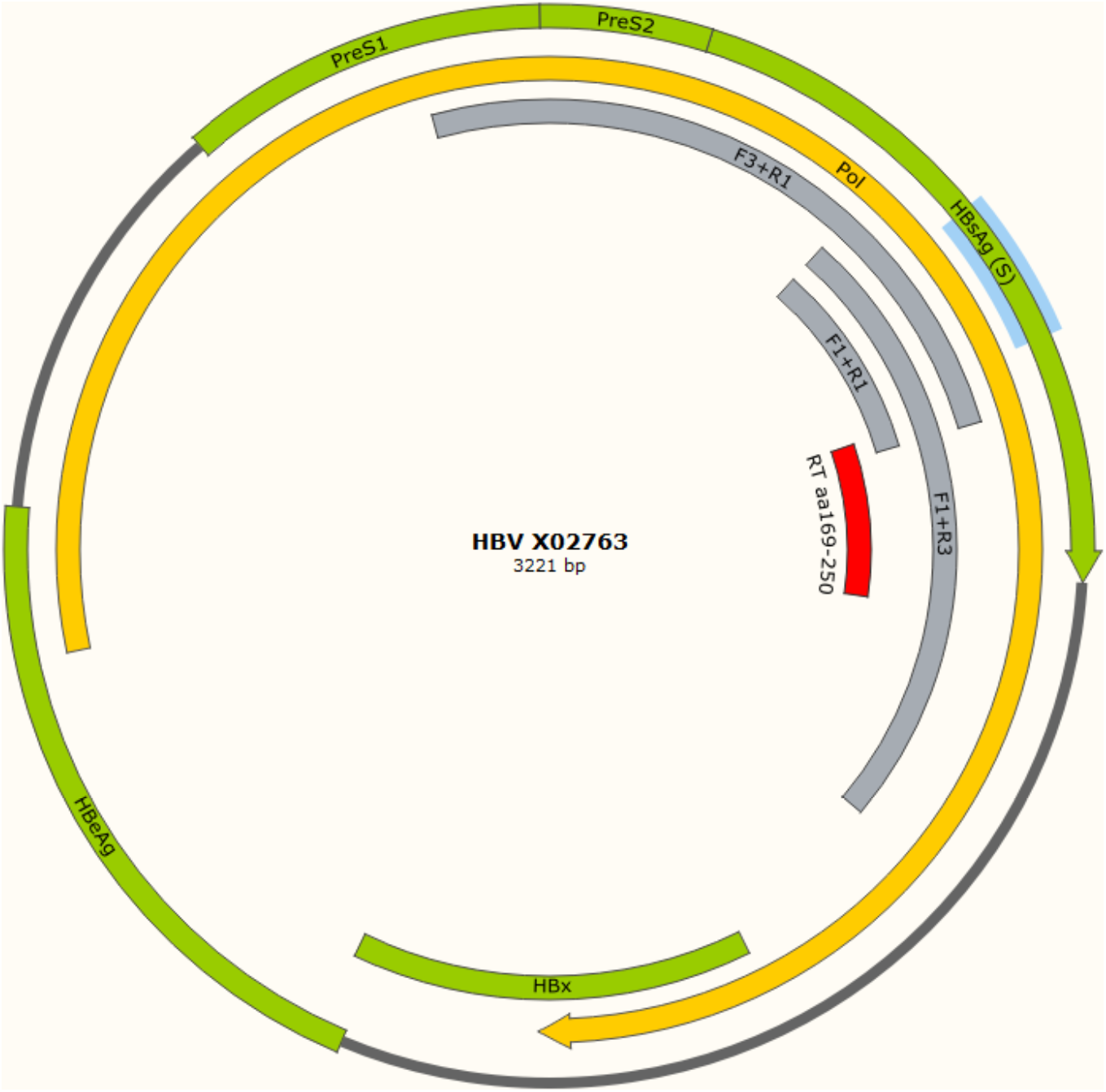
Schematic of HBV PCR amplicons produced from cccDNA. Regions covered by PCR amplicons detailed in Table 2 are shown in grey. Overlapping ORFs in the HBV genome (genotype A reference isolate X02763) are shown in green or yellow (polymerase gene). Genome is represented as linear for clarity. F1+R1 amplicon is sufficient for genotyping and limited detection of sAg escape mutants. The region of the reverse transcriptase domain (RT) in which resistance associated substitutions (RAS) arise (aa169 – 250), shown in red, is encompassed by the F1+R3 amplicon. The region in which known surface antigen (HBsAg) immune/diagnostic escape mutants arise (aa100 – 147), highlighted in blue, is covered by all three amplicons.

**Table 3:** Amplification of S/Pol gene from DSS of HBV-positive samples from Brazil, allowing genotyping and characterisation of RAS, is dependent on viral load. Clinically significant RAS in the reverse transcriptase (RT) domain of the Polymerase open reading frame (ORF) are noted. TDF, tenofovir; 3TC, lamivudine; EFV, efavirenz; nd, no treatment data available; ng, not genotyped. Blank fields for primer pair products indicate no amplicon was generated.

| Sample | Viral load (IU/mL) | Therapy | F1+R1 product | Genotype | F1+R3 product | RT domain RAS | F3+R1 product | sAg escape mutants |
| --- | --- | --- | --- | --- | --- | --- | --- | --- |
| Br1 | 816,483,772 | No treatment | Y | A1 | Y | - | Y | Y100C |
| Br2 | 389,547,123 | Peg-IFN | Y | A1 | Y | - | Y | Y100C |
| Br3 | 241,145,063 | TDF, 3TC | Y | D2 | Y | V173L, L180M M204V | Y | - |
| Br4 | 33,875,365 | TDF, 3TC, EFV | Y | A1 | Y | - | Y | Y100C |
| Br5 | 21,807,740 | No treatment | Y | D3 |  |  | Y | - |
| Br6 | 6,219,005 | TDF | Y | D3 | Y | - | Y | - |
| Br7 | 5,843,310 | No treatment | Y | A1 | Y | - | Y | Y100C |
| Br8 | 5,217,998 | nd | Y | D3 | Y | - | Y | - |
| Br9 | 3,166,098 | TDF | Y | F2 | Y | V173L |  | - |
| Br10 | 692,965 | nd | Y | A2 |  |  |  |  |
| Br11 | 193,271 | TDF | Y | F2 |  |  |  |  |
| Br12 | 94,152 | nd | Y | D3 |  |  |  |  |
| Br13 | 59,927 | TDF | Y | D3 |  |  | Y |  |
| Br14 | 43,005 | TDF, 3TC | Y | A1 |  |  |  | Y100C |
| Br15 | 35,672 | TDF | Y | A1 |  |  |  |  |
| Br16 | 25,791 | TDF | Y | D2 |  |  | Y |  |
| Br17 | 17,280 | nd | Y | F1 |  |  |  |  |
| Br18 | 16,697 | nd | Y | D3 |  |  |  |  |
| Br19 | 12,446 | 3TC | Y | D3 |  |  |  |  |
| Br20 | 12,196 | No treatment | Y | D1 |  |  |  |  |
| Br21 | 10,058 | TDF | Y | F2 |  |  |  |  |
| Br22 | 7,199 | TDF |  | ng |  |  |  |  |
| Br23 | 6,208 | No treatment | Y | D3 |  |  |  |  |
| Br24 | 4,091 | TDF | Y | B1 |  |  |  |  |
| Br25 | 2,884 | Peg-IFN | Y | A1 |  |  |  |  |
| Br26 | 2,431 | No treatment | Y | D1 |  |  |  |  |
| Br27 | 2,262 | No treatment | Y | A2 |  |  |  |  |
| Br28 | 1,069 | nd |  | ng |  |  |  |  |
| Br29 | 1,046 | No treatment |  | ng |  |  |  |  |
| Br30 | 635 | No treatment | Y | D3 |  |  |  |  |

It was not clear why three of the samples could not be amplified using the F1/R1 primer pair, but it is likely due in part to lower viral load as all three had viral loads of ~1 × 10^3^ – 1 × 10^4^ IU/mL (table 3). As the genotypes of these samples had not been previously determined it was not possible to confirm whether the primer pair had a lower sensitivity for some genotypes.

Successful amplification of the larger products using primer pairs F1/R3 and F3/R1 appeared to be associated with viral load (table 3). Both primer pairs were able to generate amplicons from samples of viral load >10^6^ IU/mL. F3/R1 displayed a higher sensitivity for genotype D samples as amplicons were achieved for samples Br13 and Br16 (viral loads 5.99 x10^5^ and 2.58 x10^5^ IU/mL, respectively), although no amplicon was obtained for the higher viral load sample Br12 (9.41 × 10^5^ IU/mL).

### Genotyping and RAS characterisation

The amplicon produced from the F1/R3 primers permitted Sanger sequencing across aa 169 to aa 250 of the reverse transcriptase (RT) domain of the Pol gene (figure 1, highlighted in red), allowing identification of resistance associated substitutions (RAS). A restricted section of this crucial RT RAS region can be sequenced with the amplicons produced from the other two primer pairs. Of those samples for which an amplicon was obtained, two samples contained known polymerase RAS. The dominant sequence amplified for Br3 contained V173L, L180M and M204V, which in isolation or combination confer resistance to lamivudine, entecavir and telbivudine. The predominant sequence in sample Br9 only contained the V173L mutation which, in isolation, is not associated directly with resistance to any of the clinically approved polymerase inhibitors.

Having refined the workflow for HBV genotyping and RAS-typing by Sanger sequencing from DSS of samples with known viral load, we applied it to a second sample set to assess the broader application of the workflow. Seventy serum samples isolated from HBV-positive patients in the Kurdistan Region of Iraq were processed onto DSS cards and shipped to the UK for analysis. Amplicons were obtained for 11 of the 70 samples and Sanger sequencing revealed they were all genotype D (table 4). Although successful amplification correlated with higher viral load (figure 2), the sensitivity of the amplification PCR for these samples was markedly lower compared to the Brazilian cohort. It is not clear why it was possible to generate amplicons for two Iraqi DSS samples of comparatively low viral load (1.15 × 10^4^ and 2.58 × 10^2^ IU/mL), or why three high viral load samples could not be amplified (2.28 × 10^8^, 1.60 × 10^7^ and 1.42 × 10^7^ IU/mL).

**Table 4.** Amplification of S/Pol gene from DSS of HBV-positive samples in Iraq cohort. Amplicons were generated from 11 of 71 DSS samples collected in the Kurdistan Region of Iraq. Clinically significant RAS in the reverse transcriptase (RT) domain of the Polymerase open reading frame (ORF) are noted.

| Sample | Viral load (IU/mL) | F1+R1 product | Genotype | F1+R3 product | RT domain RAS | F3+R1 product | sAg escape mutants |
| --- | --- | --- | --- | --- | --- | --- | --- |
| lq1 | 745,867,080 | Y | D1 | Y | - | Y | - |
| lq2 | 480,199,200 | Y | D1 | Y | - | Y | - |
| lq3 | 369,094,710 | Y | D1 | Y | - | Y | - |
| lq4 | 361,383,300 | Y | D1 | Y | - | Y | - |
| lq5 | 330,190,500 | Y | D1 | Y | - | Y | - |
| lq6 | 17,996,700 | Y | D1 | Y | - |  | P120S |
| lq7 | 12,637,111 | Y | D1 | Y | - | Y | P120S |
| lq8 | 3,393,900 | Y | D3 | Y | - |  |  |
| lq9 | 1,843,350 | Y | D1 | Y | - |  | G145R |
| lq10 | 11,514 | Y | D1 | Y | - |  | Q101H, Y143H |
| lq11 | 258 | Y | D1 | Y | - |  |  |

**Figure 2.**
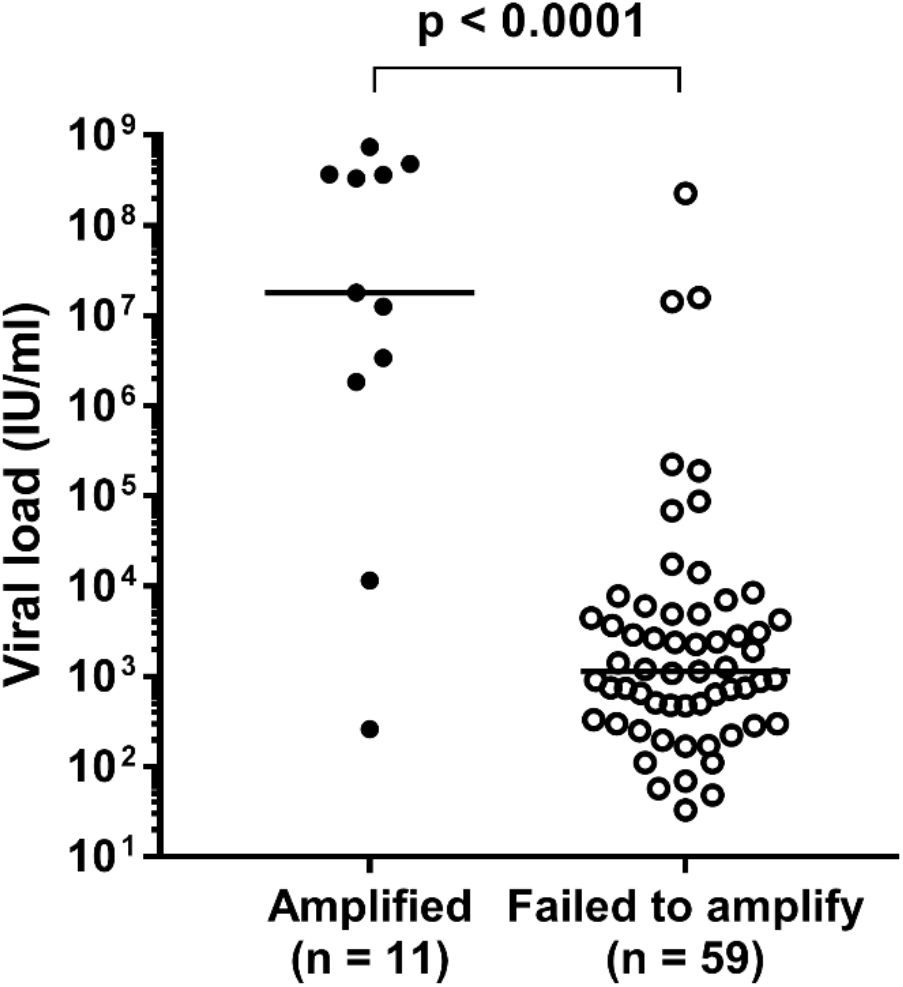
HBV viral load of DSS from Iraq. Successful PCR amplification of HBV-positive DSS samples from Iraq is associated with higher viral load (Mann-Whitney U test). Number of samples in each group (n) is shown. PCRs for samples which failed to amplify were performed at least 3 times.

While no RT RAS were detected, three samples contained known sAg immune escape mutants, demonstrating that this process can provide clinically relevant sequence information from poorly characterised samples.

Samples from both cohorts demonstrated clustering by genotype when plotted on a maximum likelihood tree with respective genotype references from GenBank (figure 3).

**Figure 3:**
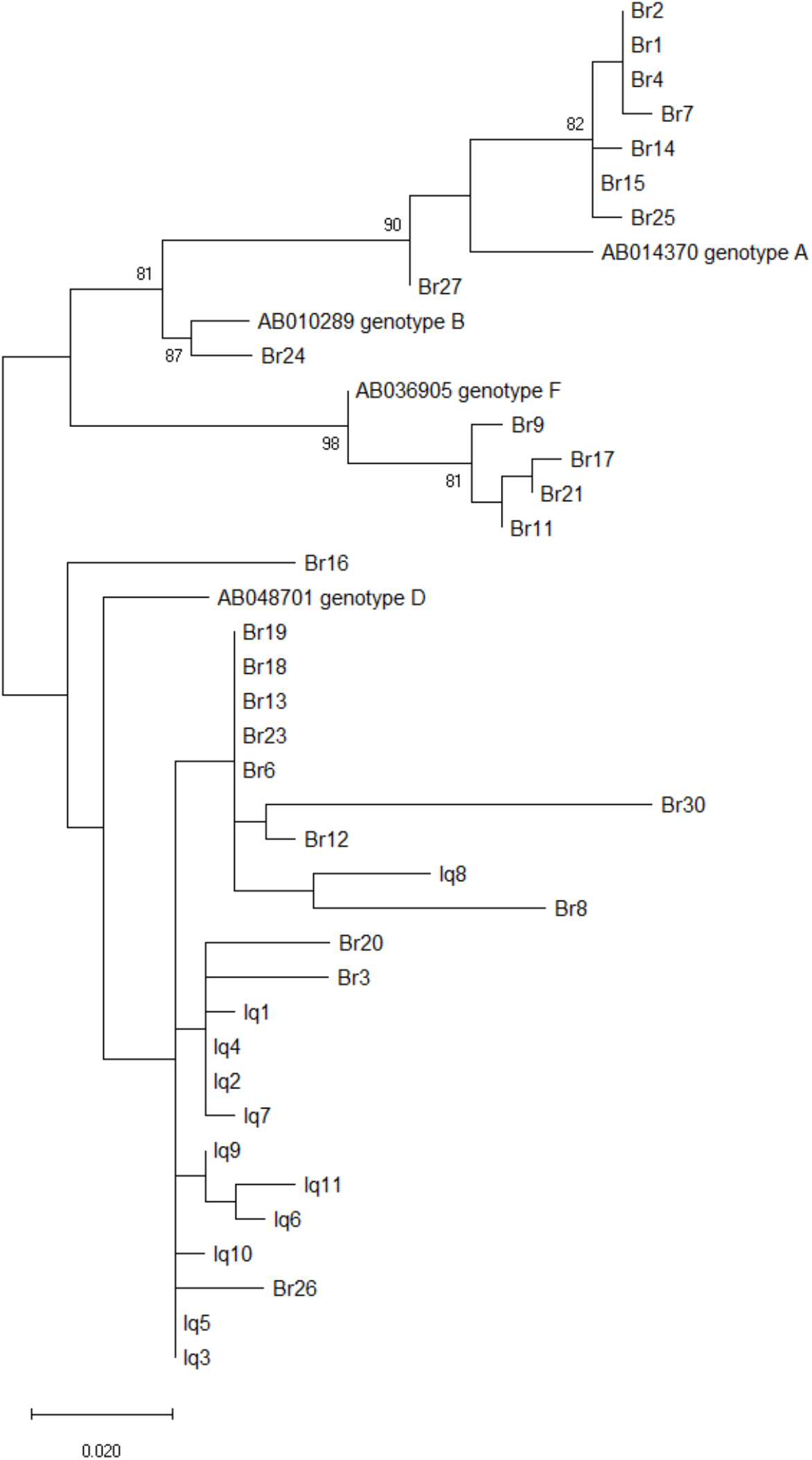
Maximum likelihood (ML) tree constructed from F1+R1 PCR amplicon (240bp). The ML tree was inferred using a general time reversible model within MEGA X (11). Statistical robustness was assessed using bootstrap resampling of 1,000 pseudoreplicates, and the tree with the highest log-likelihood is shown.

### MinION sequencing direct from DSS

Having obtained clinically relevant sequence data by Sanger sequencing from DSS, we investigated whether comparable data could be obtained using MinION sequencing. Three samples each were selected from the Brazilian and Iraqi cohorts (Br1, Br2 and Br3; Iq2, Iq3 and Iq4), that had been successfully amplified with all three primer pairs. Amplicons of 1,274 bp were generated using primers F4 and R4, which were analysed in parallel by Nanopore sequencing. A total of 185,349 raw reads were obtained for the 6 samples, ranging from 23869 to 41392 per barcode. Following adapter trimming and further filtering of erroneously long and short reads these counts ranged from 7676 to 11832 per barcode (table 5). A summary of raw sequencing reads acquired over time, and average quality score per read over time is included in figure 4.

**Table 5.** MinION sequencing yields. Raw reads are those assigned barcodes by Albacore before any further quality control. Adapter trimmed reads are those exceeding a mean Phred score of 7 and processed by Porechop. Length filtered reads were processed by NanoFilt.

| Sample | Raw reads | Adapter trimming | Length filtered |
| --- | --- | --- | --- |
| Br1 | 24105 | 13320 | 11272 |
| Br2 | 23868 | 12640 | 10309 |
| Br3 | 41392 | 14702 | 11832 |
| lq2 | 27860 | 11654 | 9572 |
| lq3 | 27478 | 12445 | 7676 |
| lq4 | 40646 | 13539 | 8979 |

**Figure 4.**
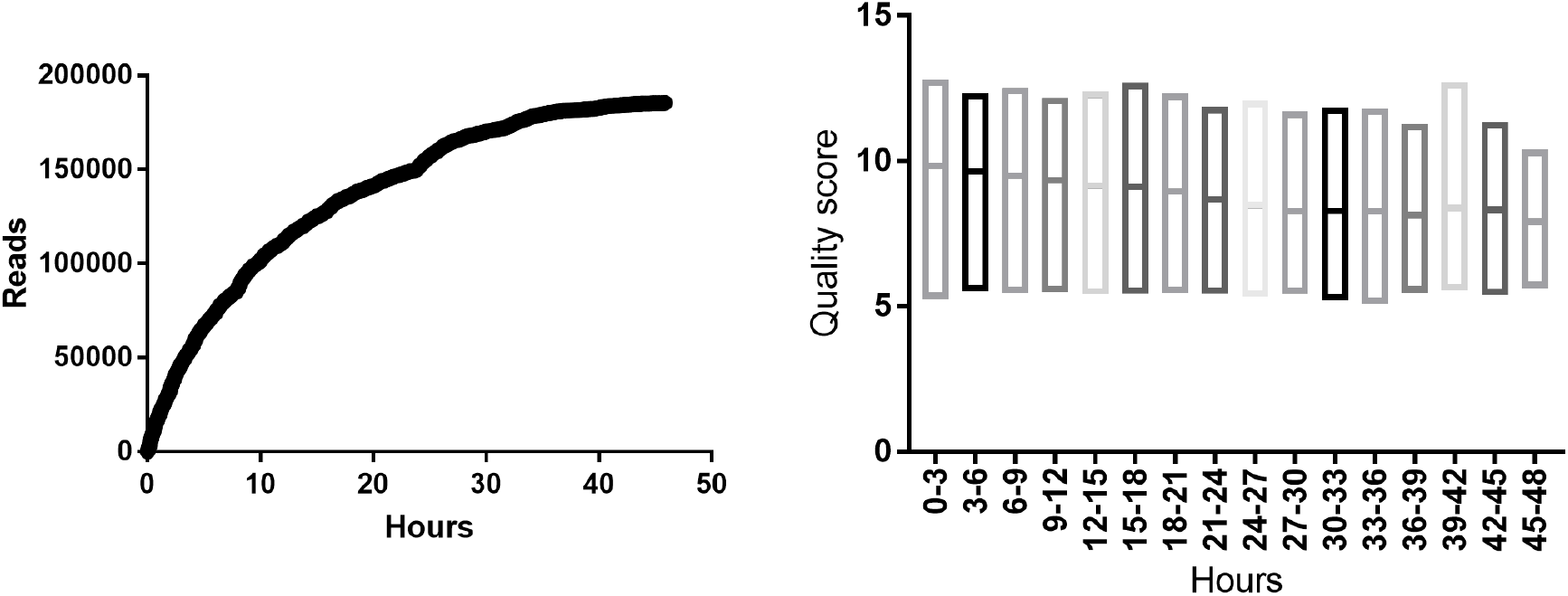
Metrics from MinION sequencing run. Both plots were generated from raw reads assigned barcodes by Albacore without further filtering. Quality scores are standard Phred scores produced by Albacore during basecalling, data is presented as mean Phred score per read with min and max.

#### Accurate consensus sequences generated by MinION Nanopore sequencing

Consensus sequences were assembled using Canu. Following assembly, the only errors observed were single base deletions within homopolymers. These ranged from 5 to 12 deletions within the total amplicon (table 6). Following a single round of Nanopolish all sequences were identical to their Sanger counterparts. Workflows for the two sequencing procedures are shown in figure 5. Upon receipt of samples it is feasible to obtain accurate sequences either by Sanger or MinION within 1-2 days.

**Table 6:**
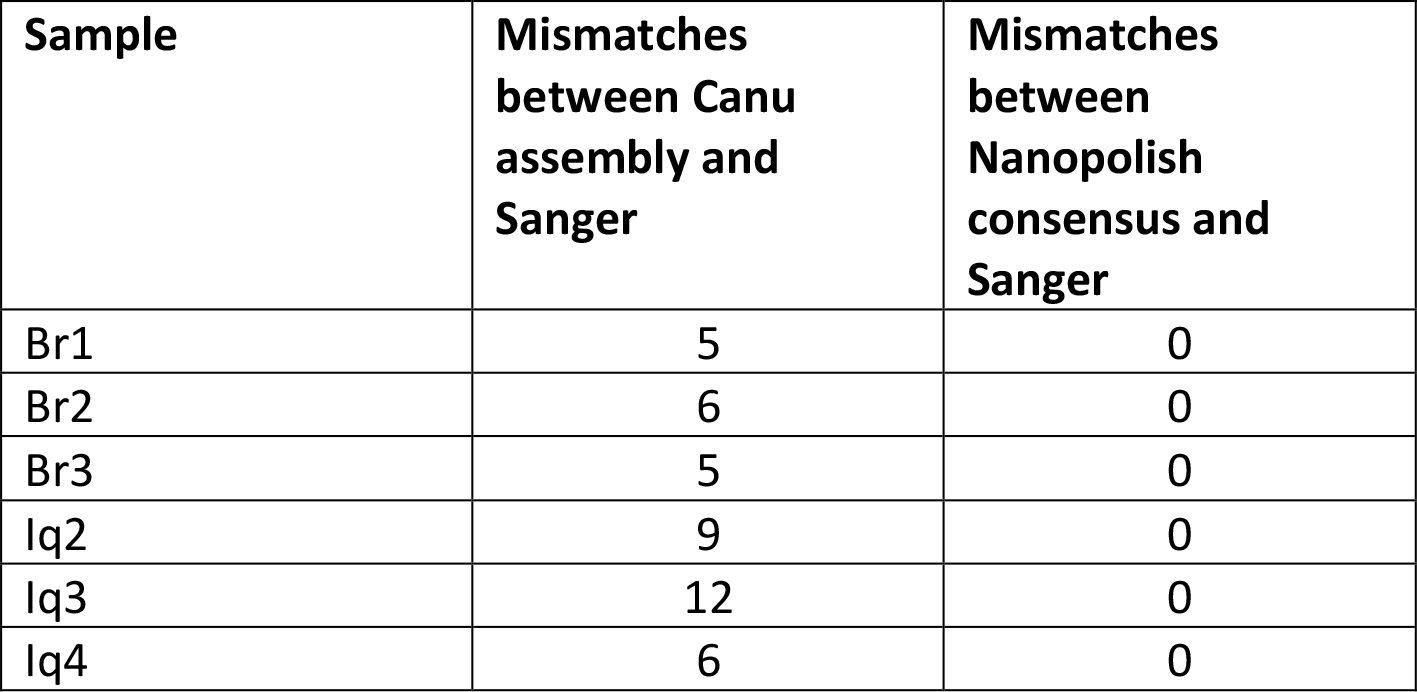
Agreement between minION sequenced samples and Sanger across the 1,274bp region studied.

**Figure 5.**
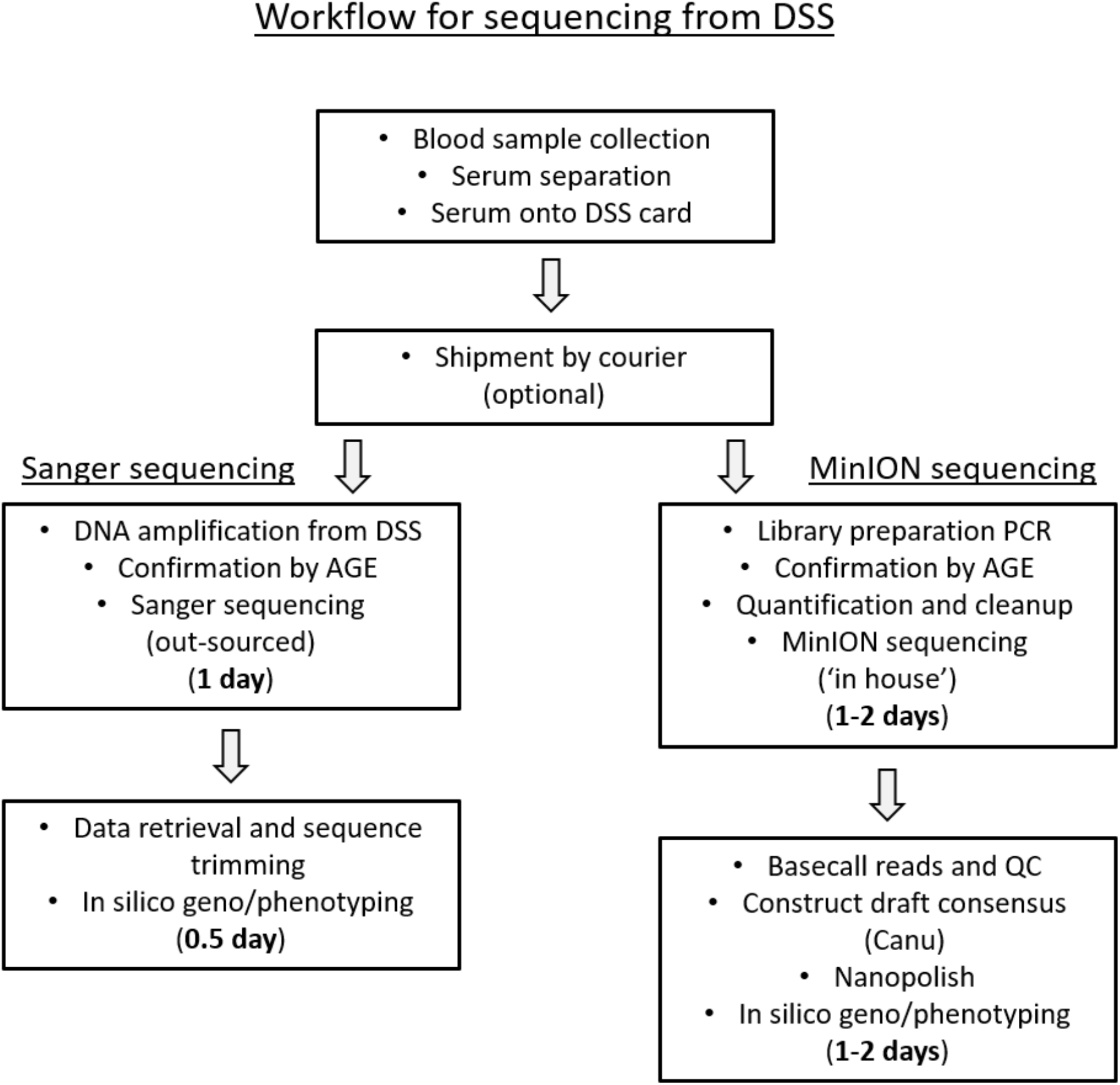
Schematic of the sequencing workflows. The typical number of days taken for each aspect of the workflow is noted. The length of time given to the ‘shipment by courier’ and Sanger sequencing steps are relevant to this study but may be omitted, or vary significantly, depending upon the given laboratory and infrastructure situation. 1 day is suggested as the minimum amount of time for MinION library preparation, sample clean up and enough sequencing time to generate a reliable consensus. Sequencing to a greater depth will add more sequencing and subsequent compute time. Sanger sequencing in Nottingham is provided by a commercial service with sample collection and over-night sequencing and data return. DSS, dried serum spot; AGE, agarose gel electrophoresis; QC, quality control.

#### Detection of minor variants with minION

A number of putative minor variants were detected by aligning minION reads with the consensus sequence, the majority of which were filtered out by Nanopolish. Of the remaining variants samples Br1, Br2, Iq1, Iq3 and Iq4 contained sequences with the P217L polymorphism in sAg in around 18% of reads, which is associated with immune escape. Iq3 also contained a L109P mutant in sAg in 16% of sequencing reads, associated with increased hydrophobicity. However, comparison with Sanger sequencing showed that neither of these base changes were visible as minor peaks on the electropherogram of the corresponding samples.

## Discussion

HBV has the capacity to exist in a host as a population of closely related viruses (18). While not displaying the extreme variability associated with RNA viruses, error-prone reverse transcription facilitates rapid adaptation to the host environment. As a consequence, resistance to antiviral therapies is common. Methods to both genotype and characterise RAS in parts of the world without access to conventional laboratory and sequencing facilities would be clinically beneficial. These techniques may become increasingly important in places such as sub-Saharan Africa, where antiretroviral programmes are in place for HIV treatment but HBV prevalence and development of resistance to drugs such as lamivudine is not accounted for (19, 20). As HIV mortality decreases and life expectancy increases, the development of chronic liver disease due to HBV infection will be of increasing concern.

We have demonstrated that clinically useful HBV sequence data can be obtained from DSS, with a lower-limit of detection of approximately 600 IU/mL, using a relatively cost-effective method that requires no cold chain. Several of the RAS detected in this cohort are clinically significant. The Y100C mutant in the S region of HBsAg has been linked to false negative/low HBsAg tests (21), and appears to be common in our Brazilian cohort, being present in 5/10 genotype A samples. More significantly, as it targets the RT domain of Pol, this method allows characterisation of drug resistance markers, which is potentially valuable in countries where prescription of drugs known to select for HBV resistance is common. The V173L, L180M and M204V RAS detected in the polymerase gene indicate resistance to the nucleoside analogues lamivudine and entecavir.

Although our original methodology removed the need for a cold chain and required only basic or mobile laboratory facilities, a Sanger sequencer (or access to a sequencer-equipped laboratory to where DSS/amplicons can be shipped) was still required. We therefore aimed to combine our DSS method with nanopore sequencing, increasing portability and to provide a method that can be rapidly deployed in resource-poor settings.

NGS datasets were successfully generated for all of the six of the samples tested. All samples demonstrated the characteristic error profile associated with nanopore sequencing, with frequent indels in homopolymers of 3 bases or more (22). In all cases these miscalls were removed using Nanopolish. These errors, which occur when voltage signal remains constant across homopolymer templates passing through the nanopore, will potentially become less of an issue with improved software interpolation or the development of new protein nanopores. In all cases the data was sufficiently accurate that the assembly generated by Canu, including indels, could be used to reliably and quickly genotype without any further processing, and with subsequent processing in Nanopolish produced sequences that were 100% identical to the sequence reads generated by Sanger sequencing. This method is therefore of use in situations where Sanger sequencing is unavailable and illustrates the utility of the MinION as a portable sequencer to determine the consensus of a virus population.

Sanger sequencing of HBV enables characterisation of a consensus sequence of theoretically the most abundant nucleotide at each position in a viral sequence within a sample.

Inspection of minor peaks on the electropherogram allows detection of minor sequence variants, down to a prevalence of around 20%, dependent on sample quality. As has been previously shown with HBV, next generation sequencing approaches allow characterisation of minor variants at a lower threshold (23-25). This information is potentially clinically useful, enabling earlier detection of resistant viruses, informing selection of appropriate treatment regimens. Accurately determining these minor variants with NGS is methodologically difficult, as low concentrations of viral template often necessitate the use of PCR before sequencing. This may introduce bias from a range of sources. A recent investigation (26) suggested that if PCR primers are well designed an amplicon sequencing approach can be as reliable as unbiased metagenomic sequencing. However, the authors highlighted that in its current level of development, MinION sequencing is not appropriate for discovering novel, low-level intra-host sequence variants due to the high false-discovery rate. In our data, we applied a conservative variant calling approach by aligning all sequence reads to the consensus followed by variant calling using Nanopolish.

Sequencing HIV from dried blood spots using 454 pyrosequencing previously showed almost complete concordance with Sanger (27), but discrepancies appeared when characterising minor variants, particularly in low titre samples (28). Further experiments using more appropriate variant calling workflows, and a side-by-side comparison of DSS, direct serum and DNA extract would be required to study the potential effect of storing DSS samples on measures of diversity. The majority of candidate variants in our data were filtered out using Nanopolish, suggesting they could be false-positives caused by sequencer error. A variant that remained unfiltered was present in 5 of 6 samples (and therefore was not genotype specific), the mutation, P217L in S has previously been linked to diagnostic escape (29). The other minor variant, L109P in S, was present in a single sample (Iq1), and has been linked to an increase in hydrophobicity (30), but the significance of this phenotype has not been determined.

The MinION sequencing protocol described here followed the manufacturer’s recommended protocol, using the standard 48-hour script in order to maximise read depth. As demonstrated elsewhere (31), an alternative approach is to run the MinION for a shorter amount of time, flush the flow cell and add different samples. Larger barcode kits (up to 96) and custom barcodes can also be used. With this assay, setup can be adjusted to favour either assay throughput or read depth, prioritizing rapid consensus sequence generation or interrogation of intra-sample diversity. The metrics produced by the sequencer as shown in figure 4 can aid in this optimisation and be used to determine the optimum timepoint to stop a sequencing run and add more samples, in order to ensure efficient use of resources. Based on current prices, and after an initial outlay on basic lab equipment (PCR machine, pipettes), we estimate that accurate sequence data with the MinION could be obtained for as little as 20 GBP per sample. In comparison, a single Sanger read across the region studied in this paper will cost ~4 GBP, with larger regions requiring at least 2 reads for complete coverage.

One limitation of this method is the amount of template input required for successful amplification; obtaining long amplicons (in this case ~1247 bp) suitable for MinION sequencing required serum spots generated from samples with viral titre of at least 10^6^-10^7^ IU/mL. As expected, successful amplification was inversely correlated with viral load, making assays which involve amplifying all or most of S in a single amplicon only applicable to high titre samples. The combination of F4/R4 primers was chosen to take advantage of the long reads available with MinION sequencing. Increased sensitivity could be achieved using a PCR-tiling approach of two or more amplicons that are pooled before sequencing (as demonstrated with the much larger Zika genome (7)), or by scaling up reaction mixtures and using more DSS per reaction.

The workflow described in figure 6 is intended as a starting point for a diagnostic workflow, but, as indicated above, this method can be modified at each point to address specific clinical questions with different pathogens. In our application, dried serum spots were studied based on available samples, but the modified *Taq* enzymes used are suitable for use with whole blood, plasma or serum added directly to the PCR reaction. There is also the potential to apply this approach to other viruses where high-throughput testing is desired, for example improving existing workflows for cytomegalovirus (32).

In summary, we describe two approaches for rapid genotyping and RAS detection in hepatitis B virus, which are applicable in resource-limited settings and require little existing infrastructure. The results presented here demonstrate the utility of direct PCR enzymes and DSS together in a clinical context. To our knowledge this has not been previously described. We have also demonstrated, for the first time, that nanopore sequencing can be applied directly to samples amplified from serum spots. Reliable sequence data was generated using the MinION sequencer, removing the need for reliance on laboratory infrastructure. Although accurate consensus sequences can be generated, it is likely that both the software and hardware associated with nanopore sequencing will require improvement before this method will find routine use for detecting minor variants within a virus population.

## Author contributions

CPM, AWT, WLI and SA designed the experiments; MMCNS, ACGJ, JFS, PJ, CHS and FTS provided samples and associated clinical data and prepared DSS cards; SA, EP and CPM carried out the experimental work; SA carried out nanopore sequencing and associated bioinformatics; SA, BK, AWT and CPM analysed the data and drafted the manuscript; all authors edited the manuscript.

## Supplementary data

Bioinformatics workflow

Following basecalling using Albacore, the “pass” folder (reads exceeding a mean Phred score of 7) was used as the input for Porechop as follows:

*porechop -i source_directory --discard_middle --require_two_barcodes*

Filtering based on length was carried out in NanoFilt:

*cat reads.fastq | nanofilt -l 1200 --maxlength 1300 > reads.filtered.fastq*

Downsampled, filtered FASTQ files for each sample were then used as the input for Canu:

*./canu --nanopore-raw reads.filtered.fastq genomeSize=1300 stopOnReadQuality=false -d canu_out -p sample-ID*

This generates several candidate contigs, the contig with the highest number of reads used was verified using BLAST and taken forwards to the next step.

Reads were then aligned in Minimap2 to their respective contig generated using Canu:

*minimap2 -ax map-ont canu_contig.fasta reads.filtered.fastq | samtools view -bS - | samtools sort -o sample.minimap.sorted.bam*

This alignment was then used to generate a polished consensus sequence using Nanopolish. First the reads are indexed to match every read in the.fastq file with its corresponding raw fast5 file (the original output of the minION sequencer):

*nanopolish index -d fast5_directory -s sequencing_summary.txt reads.filtered.fastq*

The alignment, reference contig and fastq for each sample were then used as the input for Nanopolish:

*nanopolish variants --consensus --fix-homopolymers -b sample.minimap.sorted.bam -g canu_contig.fasta -r reads.filtered.fastq -o sample.polished.consensus.vcf*

This consensus.vcf file was then converted to standard.fasta format:

*nanopolish vcf2fasta -g canu_contig.fasta sample.polished.consensus.vcf > sample.polished.consensus.fasta*

The output consensus sequence can then be checked against the original Canu contig, as well as a Sanger contig from the same sample if available. These sequences can also be used for genotyping and resistance typing against established reference sequences.

Intra-sample variants can be determined by aligning all reads to the Nanopolished consensus sequence or Sanger contig and using the *variants* function within Nanopolish to generate a.vcf file:

*nanopolish variants -b alignment.sorted.bam -g consensus.fasta -r sample.fastq -o variants.vcf --min-candidate-frequency 0.1 --ploidy 1*

---

**Supplementary table 1:**
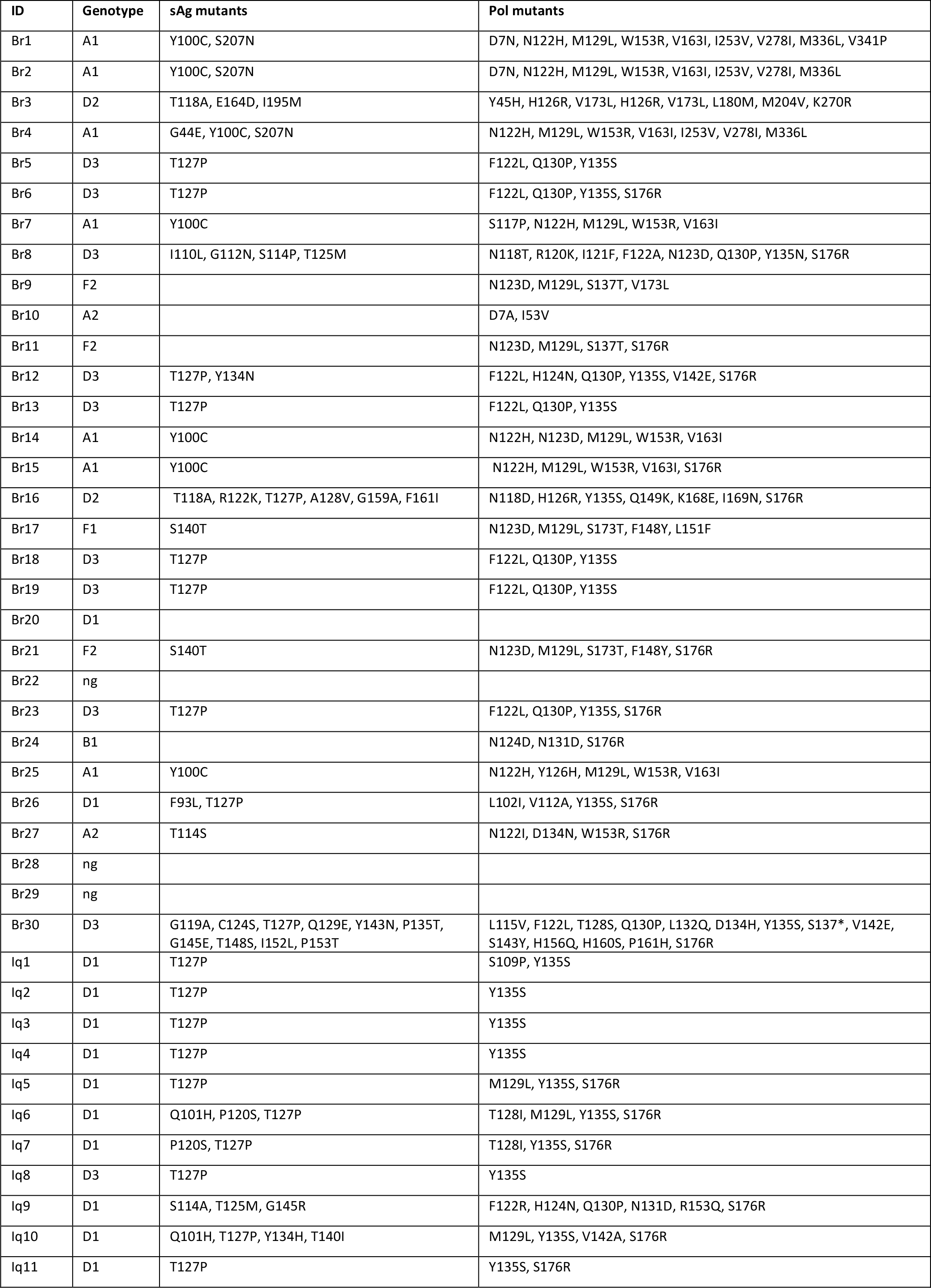
Mutations vs consensus reference sequences determined from Sanger sequencing in geno2pheno.

| ID | Genotype | sAg mutants | Pol mutants |
| --- | --- | --- | --- |
| Br1 | A1 | Y100C, S207N | D7N, N122H, M129L, W153R, V163I, I253V, V278I, M336L, V341P |
| Br2 | A1 | Y100C, S207N | D7N, N122H, M129L, W153R, V163I, I253V, V278I, M336L |
| Br3 | D2 | T118A, E164D, I195M | Y45H, H126R, V173L, H126R, V173L, L180M, M204V, K270R |
| Br4 | A1 | G44E, Y100C, S207N | N122H, M129L, W153R, V163I, I253V, V278I, M336L |
| Br5 | D3 | T127P | F122L, Q130P, Y135S |
| Br6 | D3 | T127P | F122L, Q130P, Y135S, S176R |
| Br7 | A1 | Y100C | S117P, N122H, M129L, W153R, V163I |
| Br8 | D3 | I110L, G112N, S114P, T125M | N118T, R120K, I121F, F122A, N123D, Q130P, Y135N, S176R |
| Br9 | F2 |  | N123D, M129L, S137T, V173L |
| Br10 | A2 |  | D7A, I53V |
| Br11 | F2 |  | N123D, M129L, S137T, S176R |
| Br12 | D3 | T127P, Y134N | F122L, H124N, Q130P, Y135S, V142E, S176R |
| Br13 | D3 | T127P | F122L, Q130P, Y135S |
| Br14 | A1 | Y100C | N122H, N123D, M129L, W153R, V163I |
| Br15 | A1 | Y100C | N122H, M129L, W153R, V163I, S176R |
| Br16 | D2 | T118A, R122K, T127P, A128V, G159A, F161I | N118D, H126R, Y135S, Q149K, K168E, I169N, S176R |
| Br17 | F1 | S140T | N123D, M129L, S173T, F148Y, L151F |
| Br18 | D3 | T127P | F122L, Q130P, Y135S |
| Br19 | D3 | T127P | F122L, Q130P, Y135S |
| Br20 | D1 |  |  |
| Br21 | F2 | S140T | N123D, M129L, S173T, F148Y, S176R |
| Br22 | ng |  |  |
| Br23 | D3 | T127P | F122L, Q130P, Y135S, S176R |
| Br24 | B1 |  | N124D, N131D, S176R |
| Br25 | A1 | Y100C | N122H, Y126H, M129L, W153R, V163I |
| Br26 | D1 | F93L, T127P | L102I, V112A, Y135S, S176R |
| Br27 | A2 | T114S | N122I, D134N, W153R, S176R |
| Br28 | ng |  |  |
| Br29 | ng |  |  |
| Br30 | D3 | G119A, C124S, T127P, Q129E, Y143N, P135T, G145E, T148S, I152L, P153T | L115V, F122L, T128S, Q130P, L132Q, D134H, Y135S, S137*, V142E, S143Y, H156Q, H160S, P161H, S176R |
| lq1 | D1 | T127P | S109P, Y135S |
| lq2 | D1 | T127P | Y135S |
| lq3 | D1 | T127P | Y135S |
| lq4 | D1 | T127P | Y135S |
| lq5 | D1 | T127P | M129L, Y135S, S176R |
| lq6 | D1 | Q101H, P120S, T127P | T128I, M129L, Y135S, S176R |
| lq7 | D1 | P120S, T127P | T128I, Y135S, S176R |
| lq8 | D3 | T127P | Y135S |
| lq9 | D1 | S114A, T125M, G145R | F122R, H124N, Q130P, N131D, R153Q, S176R |
| lq10 | D1 | Q101H, T127P, Y134H, T140I | M129L, Y135S, V142A, S176R |
| lq11 | D1 | T127P | Y135S, S176R |

